# Mental state space visualization for interactive modeling of personalized BCI control strategies

**DOI:** 10.1101/867119

**Authors:** Ilya Kuzovkin, Konstantin Tretyakov, Andero Uusberg, Raul Vicente

## Abstract

**Objective:** Numerous studies in the area of BCI are focused on the search for a better experimental paradigm – a set of mental actions that a user can evoke consistently and a machine can discriminate reliably. Examples of such mental activities are motor imagery, mental computations, etc. We propose a technique that instead allows the user to try different mental actions in the search for the ones that will work best.

**Approach:** The system is based on a modification of the self-organizing map (SOM) algorithm and enables interactive communication between the user and the learning system through a visualization of user’s mental state space. During the interaction with the system the user converges on the paradigm that is most efficient and intuitive for that particular user.

**Main results:** Results of the two experiments, one allowing muscular activity, another permitting mental activity only, demonstrate soundness of the proposed method and offer preliminary validation of the performance improvement over the traditional closed-loop feedback approach.

**Significance:** The proposed method allows a user to visually explore their mental state space in real time, opening new opportunities for scientific inquiry. The application of this method to the area of brain-computer interfaces enables more efficient search for the mental states that will allow a user to reliably control a BCI system.

## 1. Introduction

In many BCI experiments, participants are asked to perform certain mental actions. Consider an experiment, where a person is asked to control a point on a screen, and have it move to the left. In essense, the subject is requested to focus on a thought of ‘moving the point leftwards’. This request is quite ambiguous – should the user concentrate on the abstract notion of ‘left’, engage in motor imagery or think about an unrelated concept?

The question of which mental paradigm would be the most applicable has been evaluated in the literature. For example the work by Friedrich et al. [1] explores different mental strategies and their effect on classification performance of brain–computer interfaces (see more examples in the section on related work). In a large study by Guger et al. [2] it was observed that for a specific 2-class motor imagery paradigm out of 98 subjects 92% were able to control the system with an accuracy of 60% or higher. However, as brain activity for a particular mental action differs across subjects [3, 4, 5], any general paradigm will be suboptimal compared to a user-specific one. How would the results of that study change, if users were allowed to user personalized experimental paradigms? In this work we propose a method that facilitates self-guided interactive search for a user-specific paradigm through communication between the user and the learning system.

To achieve our goal we replace the traditional feedback [6] with a visualization of the feature space within which the underlying machine learning algorithm is operating. This visualization facilitates a ‘dialogue’ between the learning algorithm and the user by visually explaining to the user why his current set of mental actions is suboptimal, which ones are being recognized well by the system and which ones should be changed. By exploring how their mental actions affect the visualization, a user can find a suitable mental action for each of the stimuli. The exploration of the mental state space can go for as long as needed to find mental actions that the user can produce consistently over time and that are distinguishable by the algorithm.

Our method is based on a variant of a Self-Organizing Map (SOM, [7]), made to operate in an online setting and to serve as a classification algorithm in addition to its primary function of building a topology-preserving low-dimensional map of the data. We directly show that map to the user of a BCI system and allow them to observe how a 2D SOM evolves in real time while receiving new data samples from test subject’s EEG device. At the same time the underlying model is mapping the recorded signals to the desired actions and measures its classification performance.

We demonstrate the feasibility of the approach on EEG recordings from two separate experiments on muscular and mental activity. The approach is general and does not depend on the neuroimaging method.

## 2. Related work

The method we propose in our work is built on the intersection of brain-computer interface research and three other areas of scientific study: self-organizing maps (SOM) on neural data, real-time neural signal visualization, and personalization of mental strategies in a closed-loop feedback scenarios. In this section we explore the work that was done in these areas in the context of BCIs.

Since their introduction in 1990 [7] SOMs have found multiple application in the analysis of neural signals and EEG signals in particular. As a clustering method, SOM has been applied to group neural patterns according to EEG spectra [8, 9]. As a classification method, SOM was used to identify epileptic condition [10], emotions [11] and other mental states [12, 13, 14]. For brain-computer interfaces, SOMs are used to classify motor imagery [15, 16, 17] and other mental tasks [18]. Being also a dimensionality reduction technique, SOM allows to visualize the recorded neural signal for the benefit of the the user or a researcher. By visualizing a neural signal as a trajectory on a 2D SOM [19] the authors were able to analyze the variation in EEG signal and identify some overall states of the individual. This and other works [20, 21] demonstrate SOMs applicability to the analysis and characterization of brain states.

The second technological component that plays a role in our method is the idea of online visualization of a neural signal to give users real-time feedback. In [22] authors evaluate an online system that allows the user to interactively select different frequency components for different actions. In [23] the authors propose a framework for signal visualization in real time, allowing the researchers to analyze the signal while the test subject receives the feedback in a virtual reality (VR) environment. Ideas on using more engaging environments for online feedback were also proposed in [24, 25]. Some works [26] explore which features can be monitored in real time. Systems like the Mind-Mirror [27] allow the user to see their brain activity in real time, but do not provide the user with an actionable knowledge that would allow to improve the performance of the system. While these ideas explore the ways of making the feedback more informative and engaging to the user, none of them fully closes the loop by showing algorithm’s understanding of the user’s mental state space to the user directly.

Lastly, the problem of choosing the best kind of mental activity for BCI has been studied in [28, 1]. Most experiments first propose a particular paradigm and then evaluate its average effectiveness on a sample of users. Many paradigms have been evaluated this way [29, 30, 31, 32, 33, 34, 35, 36]. However, as brain activity for a particular mental action differs across subjects [3, 4, 5], any general paradigm will be suboptimal compared a user-specific one.

In this work we propose a method that facilitates self-guided interactive search for a user-specific paradigm through communication between the user and the learning system. Real-time visualization of user’s mental state representation is used directly as feedback for the user, thus allowing the user to understand and affect algorithm’s mapping of his mental states to the actions of a BCI system. This enables an active dialog between the two information processing systems: the learning algorithm and the human brain.

## 3. Methods

At the core of almost any BCI system lies a machine learning algorithm that classifies user brain signal into desired actions [37]. The algorithm sees the incoming data in a high-dimensional space and operates in that space. If an algorithm is unable to correctly discern the desired actions from the signal one can rely on visualization of the data and the space state to figure out why that is the case. Visualization allows to see particular data points in the context of other data, and allows to detect such issues as homogeneous data representation, failure to represent critical features of the data, biases in the data, insufficient flexibility of the feature space to present different data points differently, too high variance of the data points that should belong to the same group, and others. In the case of classification of mental actions we find that the two most important aspects a visualization could help evaluate are the cases where the data points from different classes look too much alike (one mental pattern is too similar to another) for the algorithm to differentiate between them, and the variance of the data within a class – mental patterns that a user produces for the same action are not consistent and the algorithm is not able to group them together. With enough simplification we were able to present such a visualization to the user directly, allowing for a real-time evaluation of the above-mentioned issues during the training process. This allows the user to modify his mental activity in accordance with the feedback and try to produce more consistent and more distinguishable mental patterns. Figure 1 depicts the interaction process between the user and the proposed feedback system.

**Figure 1.**
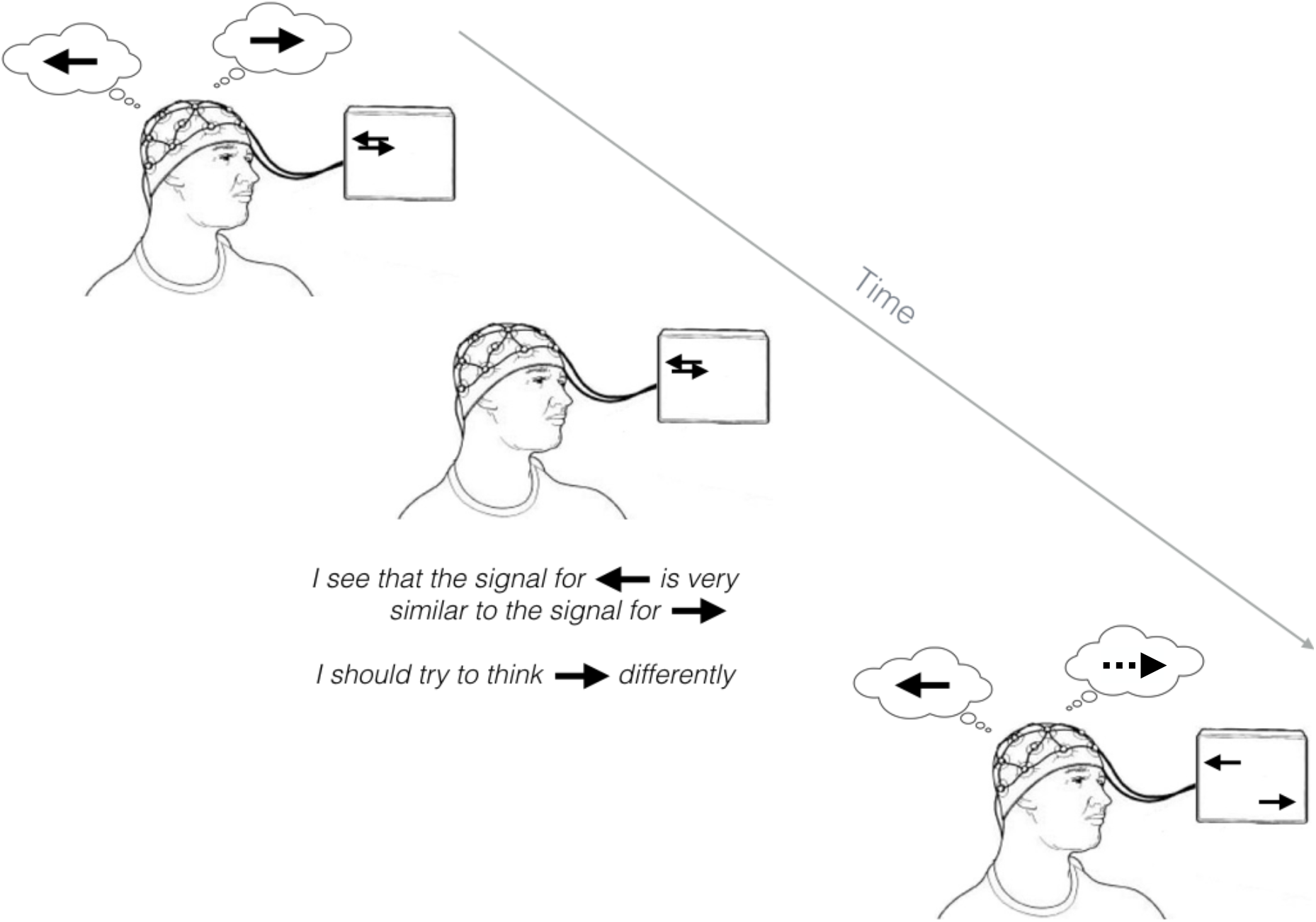
Real-time interaction process between the system and the user, during which the user realizes that he must modify his mental activity for one of the actions to increase the system’s performance.

Direct visualization of the space of mental signals provides more information to the user and allows to make more informed decisions than would be possible with the traditional approach [6]. If in the case of usual system the subject has no information of why the system cannot distinguish the user’s mental states, in the *adaptive* paradigm, proposed in this work, the subject can see which mental actions are not being recognized or are colliding with others, previously attempted, mental states. The proposed framework naturally addresses a few limitations of the traditional approach, such as limited number of actions that can be trained simultaneously and makes a more efficient use of the training time by shifting training data class distribution towards more complicated cases.

The concept described above poses several technological constraints on the choice of the underlying learning algorithm. To facilitate the real-time feedback loop the algorithm should work in an online setting and be fast enough to support real-time operation. In order to present the projection of the feature space to the user the algorithm must be compatible with topology-preserving dimensionality reduction techniques [38]. In this section we describe a method that satisfies those requirements.

### 3.1. Self-Organizing Map

Self-organizing map (SOM) [7] is one of the topology-preserving dimensionality reduction techniques. These techniques try to preserve the relative distances through the transformation, such that the data points that were close in the original space remain close in the target space, and those that were apart, remain apart. SOM projects the data form the original space onto a *map*, which is a collection of *m* units organized into a multidimensional rectangular grid. Most commonly (and also in this work) a two-dimensional grid is used. Each SOM unit *u* corresponds to a vector **w**_*u*_ ∈ ℝ^*d*^ in the space of input data points (signal samples from the EEG device, in our case). This way each unit effectively covers a region in the signal space. In this work the map has 625 units (25 × 25 square grid) with 630-dimensional vectors **w** initialized from uniform distribution 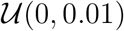.

The learning phase consists of updating vectors **w** with each new training sample **x**. Once a new sample is obtained from a neuroimaging device the *best matching unit* (BMU) for that sample is found according to Equation 1 with Euclidean distance used as the distance measure.

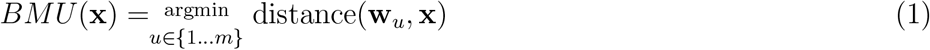

Once BMU is found the weights **w** of unit *u* and its neighbors are updated as shown in Equation 2, where *s* is the number of the current iteration.

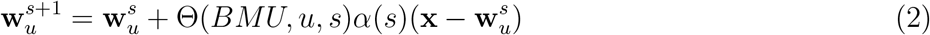

Default SOM is an offline learning algorithm that performs several passes over the training data. The update is thus repeated for each iteration *s* ∈ {1, …, *S*}, for each input data vector (**x**_1_, … , **x**_*n*_) in the training set and for each unit in the map (*u*_1_ , … , *u*_*m*_). In total this procedure is being repeated up to *S* × *n* × *m* times, where *S* is the iteration limit, *n* is the number of samples in the training data and *m* is the size of the map. Not all units are updated with each new input vector, furthermore, not all units among the updated ones are updated equally. There are two functions in Equation (2), which are responsible for deciding which units will be updated and by how much. Θ(*b, u, s*) is called the *neighborhood function*, it determines to what extent unit *u* is neighbor of a unit *b*: for *b* itself Θ(*b, b, s*) = 1 and for some unit *u*, which is too far away to be considered to be a neighbor of *b* Θ(*b, u, s*) = 0. The parameter *s* is used to decrease the number of neighbors on later iterations. The function *α*(*s*) outputs *learning rate* that decreases with more iterations allowing the learning process to converge.

At the end of the learning process the units of the map represent centers of signal clusters in the training data. Each new data sample can be assigned to one of the clusters and this cluster will hold samples that are similar. The samples that are close in the original space will be assigned to map units that are close to each other on the map.

### 3.2. Predictive Online SOM

We extend SOM to work in an online setting [39, 40], where the map is updated only once with each new data sample. We also assign a vector of probabilities **p**_*u*_ ∈ ℝ^*C*^ to each unit *u* and use that vector to classify each new incoming data sample into one of *C* classes. The class probability vector **p** of unit *u* of Predictive Online SOM (POSOM) is initialized to a random vector of length *C* with values sampled from uniform distribution 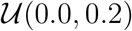. This vector holds action probability distribution for the unit *u*. It shows what is the probability that a signal **x**, which was classified into unit *u*, has been produced in response to action *a*.

Class probability vectors are updated after each sample according to Equation 3,

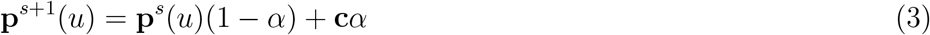

where *s* is iteration number, *α* ∈ (0, 1) is a parameter, which specifies how fast the contribution of the older data samples deteriorates, and **c** is a bit vector, where for each class we have value 0 or 1. There can be only one non-zero value in the vector **c** and its position indicates the true class of a sample.

The probability vector **p**_*u*_ is used for classification as follows: for each new sample **x** we first identify POSOM’s BMU *u* for this sample, and predict the class of this sample by choosing the most probable class in the vector **p**_*u*_.

### 3.3. POSOM-based BCI training system

The learning method defined in the previous section satisfies all of the requirements of a system with an interactive feedback based on visualization on user’s mental state space we have outlined in the introduction.

The training process begins by presenting an empty SOM map to the user (Figure 3a). A stimulus cue is displayed for a brief period of time and the system starts receiving samples from the neuroimaging device. It finds the best matching unit *u* for each of the samples and updates the **w**_*u*_ and **p**_*u*_ vectors of the unit *u* and its neighbours. Some of the randomly initialized SOM units now represent certain mental patterns and are mapped to corresponding actions, the action each unit is associated with is shown with a pictogram on the map (Figure 3b).

**Figure 2.**
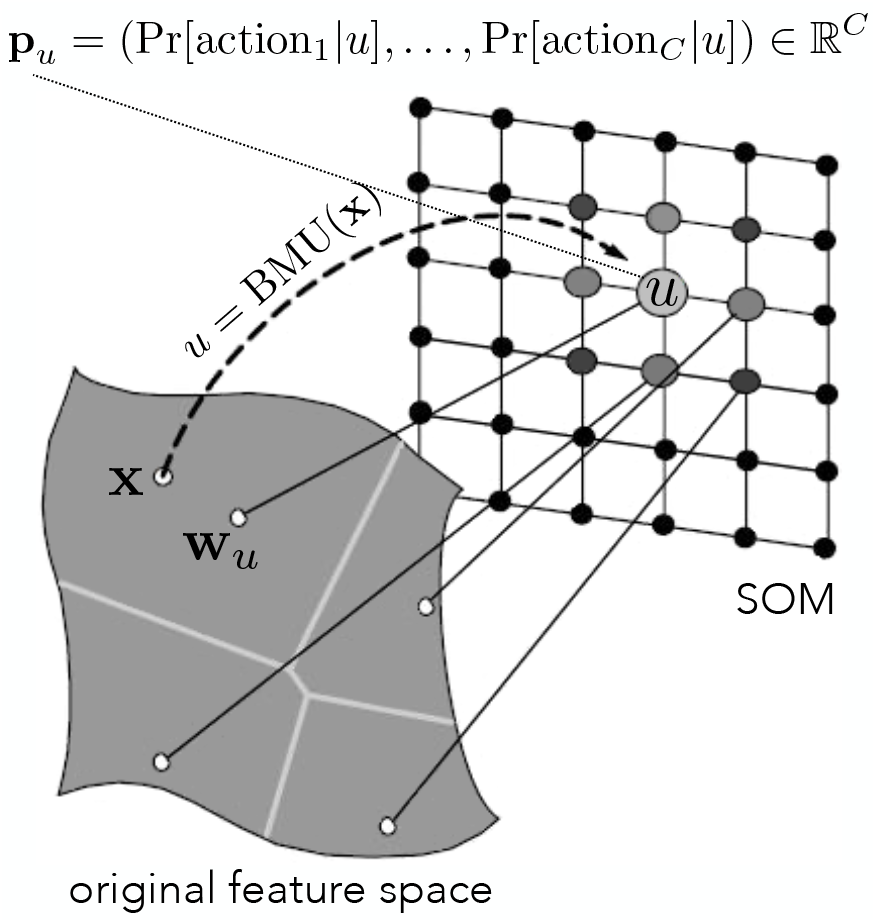
SOM extended with class probability vectors. Signal representation **w**_*u*_ in the original feature space is mapped to a unit *u* on a two dimensional map. This unit represents a cluster of signal samples similar to **w**_*u*_, such as sample **x**. Unit *u* holds a vector of class probabilities **p**_*u*_ that shows to which class a sample assigned to the cluster with centroid *u* most probably belongs.

**Figure 3.**
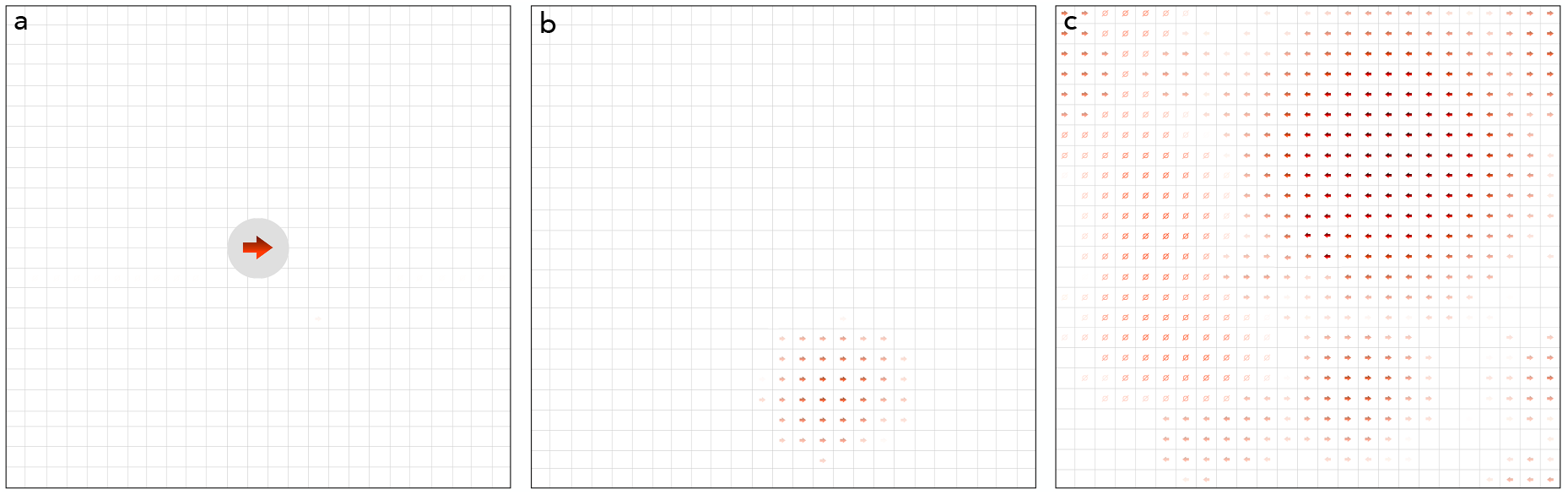
The process of forming the map. The three panels represent the visualization at three different stages. **a:** The process starts with a visualization of an empty SOM and the very first stimulus cue, in this case – the “right” action. **b:** The first few samples of a test subject thinking “right” are now collected and assigned to the units on the map, forming a region on the map that is dedicated to signals that correspond to the mental pattern “right”. **c:** Repeating steps (a) and (b) for all stimuli results in a map, where units are assigned mental state representations and corresponding actions. Bottom-right covers the “right” action, area in the top right is dedicated to the “left” action, and the left side of the map is now associated with the “relax” action. This is an example of a successful session, where a user has achieved good separation between the signals.

The process continues until the user is satisfied with his ability to produce the required set of patterns consistently and system’s ability to assign these patterns to correct units on the map (Figure 3c). The user can see on the map which of the mental patterns are always assigned correctly and which ones are ‘jumping’ across the map. This informs the user about the variance of a mental pattern, if the variance is too high it might be best to switch to another mental pattern instead. If a user can see that the patterns of two or more different actions tend to occupy the same region of the map he can conclude that the mental patterns he is producing for these actions are not different enough to be distinguishable and he should consider replacing one or all of them.

### 3.4. Experimental setup

An experimental session consists of a set of four recordings to compare maximal achievable classification accuracy of *adaptive* (proposed) versus *control* approach. Two recordings for collecting the training data for each approach and two for testing each approach. In the first pair of the experiments the test subjects were allowed to engage facial muscles in response to the stimuli, while in the second experiment they had to only use the mental activity and fully avoid facial or eye movements. We refer to these experimental conditions as *facial* and *mental* correspondingly. This first experiment was set in place to use a signal with high signal to noise ratio to validate the methodology and the second to evaluate it on a more challenging task. During the facial experiment the test subjects were instructed to consistently use any facial expressions they can think in response to the stimuli, and were presented with a list of common potential actions they could do, such as eye blinks, clenching of the teeth, smiling, etc. For the mental experiment test subjects were instructed not to move any muscles during the signal recording intervals. They were advised on the major mental activities used in BCI, such as motor imagery, mental computational, etc. In total each user completed 8 recording sessions guided by the interface depicted on figure 4: control training (4b) and testing (4c) using facial muscles as responses, adaptive training (4a) and testing (4c) using facial muscles, control training (4b) and testing (4c) using mental responses only, and adaptive training (4a) and testing (4c) using mental responses.

**Figure 4.**
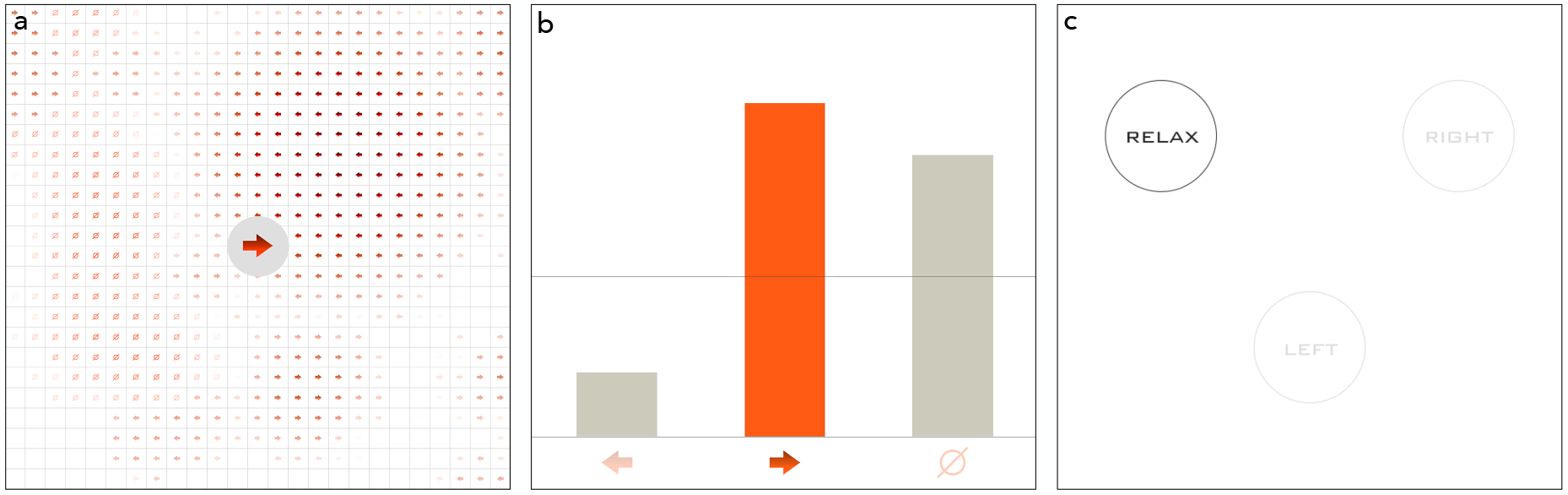
Interface of the experiment. **a:** In **adaptive** experiment the user is presented with a grid that visualizes 2D projection of decoder’s feature space. The grid is updated with each new data sample received from the neuroimaging device, enabling the user to see how his mental actions affect the mental state space representation in real time. Cue to the next stimulus is shown in the center of the screen and disappears after 1 second, allowing the user to see the full grid. **b:** The **control** experiment provides the feedback by raising or lowering per-class performance bar, indicating which stimuli are being recognized well by the system. **c:** The decoding models resulting from both adaptive and control experiments are **tested** with the same interface, where a user is presented with a description of the mental activity he must engage in. We do not use the same cues as during the training to measure the ability of the user to engage the metal activity associated with the action and avoid measuring involuntary reaction to the cue image.

The sequence of the recordings per test subject per experimental condition is as follows:

i. Training of the classification model in the traditional way. Stimuli were presented in a random order for 7 seconds each, for total time of 7 minutes, keeping the number of times each stimulus is shown balanced. The test subject received real-time feedback by observing the height of the performance indicator that was changing with each new data sample. Currently highlighted bar is the current action, height of the bar indicates the performance (Figure 4b).
ii. Testing the traditional model. To avoid measuring the involuntary reaction to the cue image the user interface of the testing stage was different from the training stage and is shown on Figure 4c. Currently highlighted stimulus is the one the user should engage in. Stimuli were shown for 7 seconds in random order for a total length of 3 minutes.
iii. Training of the classification model in adaptive way. The user was presented with a visualization of the projection of the feature space onto 2D grid (Figure 4a). Each stimulus is shown for 7 seconds, the duration of the experiment was not fixed to allow the subject to test different mental activities for the same action until the one that works is found. The stimuli were presented in the order of their performance rate, the actions that have the lowest score are shown more frequently.
iv. Testing of the adaptively trained model. The procedure repeats the steps outlined in (2) exactly, making the testing runs comparable.

Subjects were seated in front of a computer screen in a quiet room with no distractions. In total 5 subjects have participated in the study (4 males, 1 female, average age of 25.3 ± 6.0). All subjects had normal or corrected to normal vision. Informed written consent was obtained from all participants of the study. In all of experiments subjects were presented with 3 different stimuli (`left, right` and `none`) and were asked to engage different mental (or, in the case of the experiment where facial expressions were allowed, muscular) activity for each stimulus. A stimulus was shown for 7 seconds. Subjects were briefed on the usual mental paradigms including motor imagery [6, 41], vivid visual imagery [42, 43], mental computations [35] and auditory recollections [44].

To evaluate the classification accuracy of the system we used averaged per-class F1 score, a harmonic mean of precision and recall of a statistical model, computed as 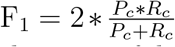 for each class *c* and averaged over all classes. Per-class precision *P*_*c*_ shows the ratio of data samples that were *correctly* identified as class *c* to all samples identified as class *c*. Per-class recall *R*_*c*_ shows the ratio of data samples correctly identified to belong to class *c* to the *total number* of samples that belong to class *c*.

Upon finishing the trials, in addition to the quantitative results, the test subjects were asked of their subjective evaluation of the adaptive system in comparison with the traditional one and whether they were able to feel the interaction with the system and its efforts to adapt to test subject’s mental efforts.

### 3.5. Preprocessing of EEG data

The data was recorded using the Emotiv EPOC [45] consumer-grade EEG device. Signal from all 14 EEG channels was split into 1000 ms windows with a step size of 250 ms. Each 1000 ms recording was linearly detrended and converted to frequency domain using fast Fourier transform [46]. Frequencies outside the 1 to 45 Hz range were excluded from further analysis.

A 1000 ms signal from one channel was represented by 45 frequency power measurements. By concatenating representations of all 14 channels we obtained feature representation of a signal with 630 dimensions. In machine learning terms a *sample* **x** that represents 1000 ms of EEG recording has 630 *features* and a categorical class label.

### 3.6. Code and data availability

The data supporting the findings of this work along with the code of the analysis and experiments are publicly available at https://bitbucket.org/ilyakuzovkin/adaptive-interactive-bci.

## 4. Results

We have conducted two types of experiments to empirically validate the benefits of the adaptive search for mental BCI paradigm via visual exploration of a projection of subject’s mental state space. During the first experiment the test subjects were allowed to engage in facial muscle activity in response to the stimuli [47, 48]. The second experiment was aimed at controlling the system via mental efforts only. In both experiments the proposed approach demonstrated statistically significant improvement in performance over the traditional method. We would like to note, however, low F1 scores obtained in the second (mental only) experiment due to the more challenging nature of the experiment and technical properties of the equipment. The results obtained in the first experiment clearly demonstrate the benefits of the proposed approach, while the results of the second (mental) experiments are to be considered preliminary, although numerically they satisfy the requirements of statistical significance. Average performance of the model trained on facial expressions in adaptive way was 23% higher (Mann-Whitney U test *p* = 0.006) than that of the traditional approach. For the mental actions the adaptive approach resulted in a model that was significantly higher than the chance level (F1 score = 0.422), while the traditional approach failed to deliver a non-trivial model (F1 score = 0.354). Comparatively, the adaptive approach yielded 19% higher performance (Mann-Whitney U test *p* = 0.018).

Figure 6 presents the detailed analysis of the results of the experiments involving facial expressions. Face muscle activity highly affects the EEG readings [49] and can be observed with the naked eye even on the raw signal. The primary goal of this series of experiments was to demonstrate the benefit of the adaptive approach in a high signal to noise ratio (SNR) scenario. During the process of interaction with the adaptive system test subject mostly converged to using such facial responses and clenching teeth, blinking one or both eyes, smiling, opening mouth and moving eyebrows. In their subjective evaluations the test subjects reported such observations as *“I was able to see that left and right eye blinks were not different enough, so I kept left blink for the left action and started opening my mouth for the right action”*, *“I tried various facial expressions but they did not work, then I tried the suggested actions and saw the difference”*, *“I felt that changing my activity affects the visualization, I was able to see how it changes”*.

**Figure 5.**
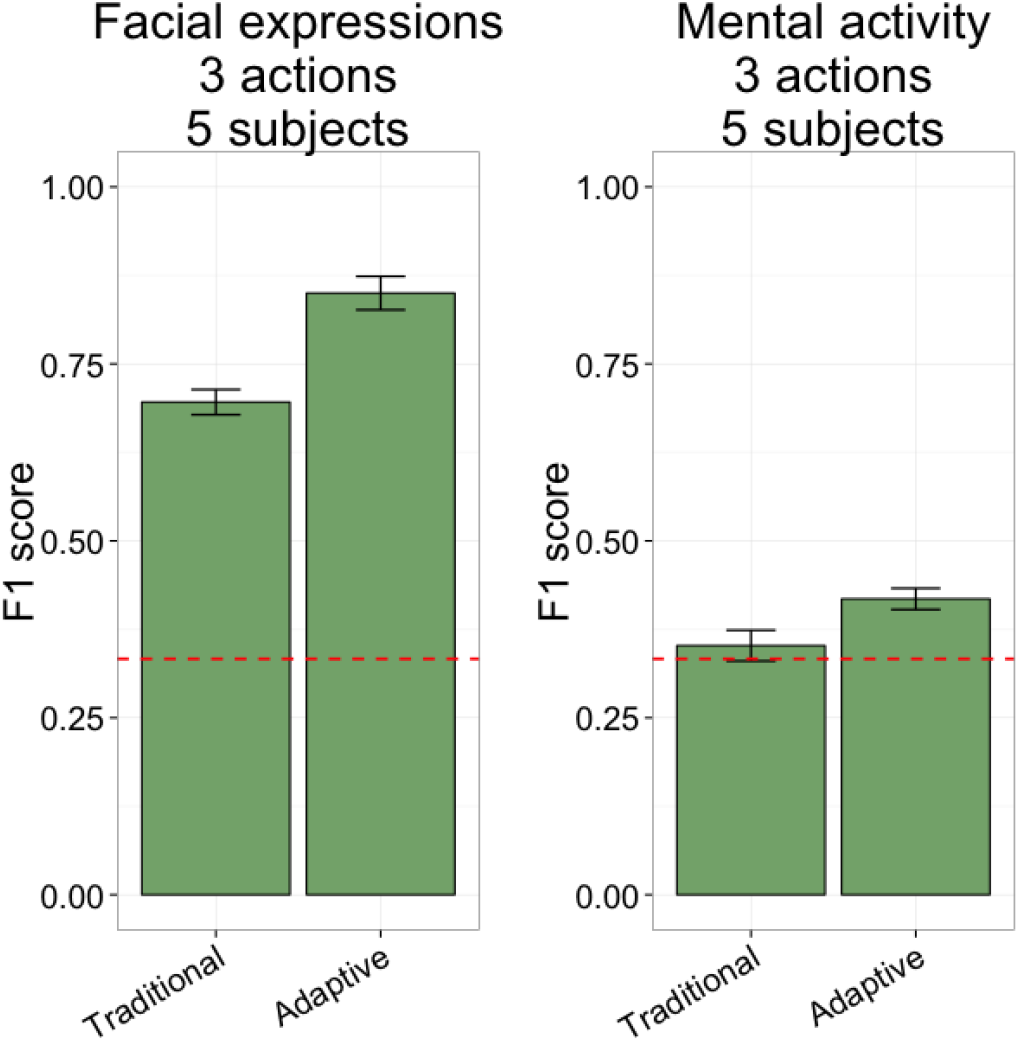
Average result of the experiments on facial expressions (left) and mental activity (right). In both cases adaptive approach demonstrates statistically significant improvement over the traditional approach.

**Figure 6.**
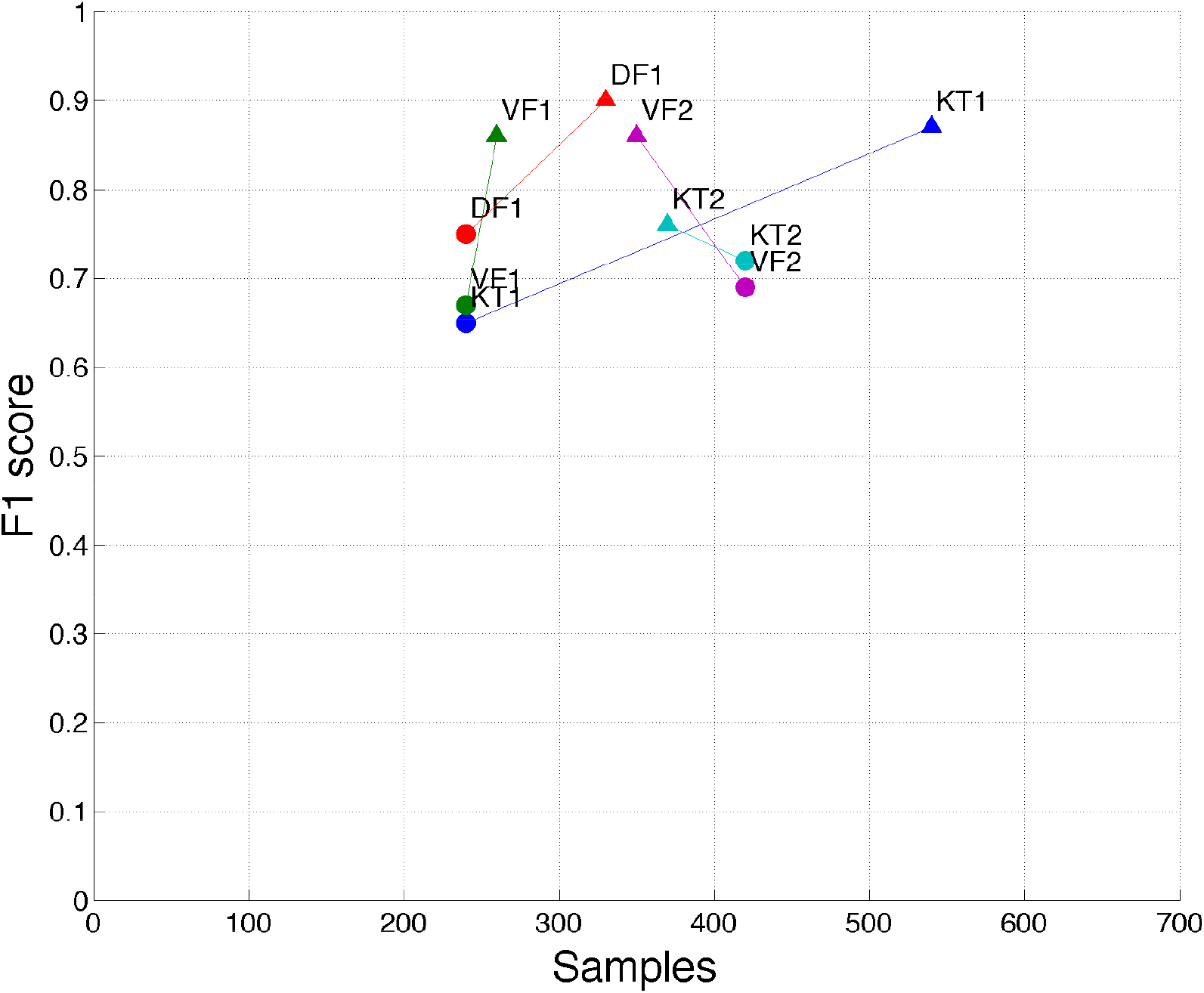
3-class training results using facial expressions. Circular markers denote the results achieved using the traditional approach and the triangular ones denote the adaptive approach. Each test subject is marked with a different color. On the *x*-axis we can see the number of samples the algorithm needed to reach the F1 score displayed on the *y*-axis. Traditional experiments were ran for 240 samples, or, if a subject felt that he would benefit from longer interaction with the system, the experiment was extended to 420 samples.

Compared to the facial expressions experiment, the task of distinguishing mental states was much harder [50]. Since the effect of changing the activity was not immediately evident, it required more time for the test subjects to begin to understand how their efforts affect the learning system. Figure 7 shows the detailed results of the experiment. The subjective reports stated that *“It was much harder”*, *“In control experiment I had no idea why it does not work, with visualization I at least saw that all my thoughts appear the same on the screen, not like it was with faces, at least I knew why it does not work”*, *“I think I was able to see an improvement when I started to relax for the neutral action and to feel angry and agitated for left and right actions”*.

**Figure 7.**
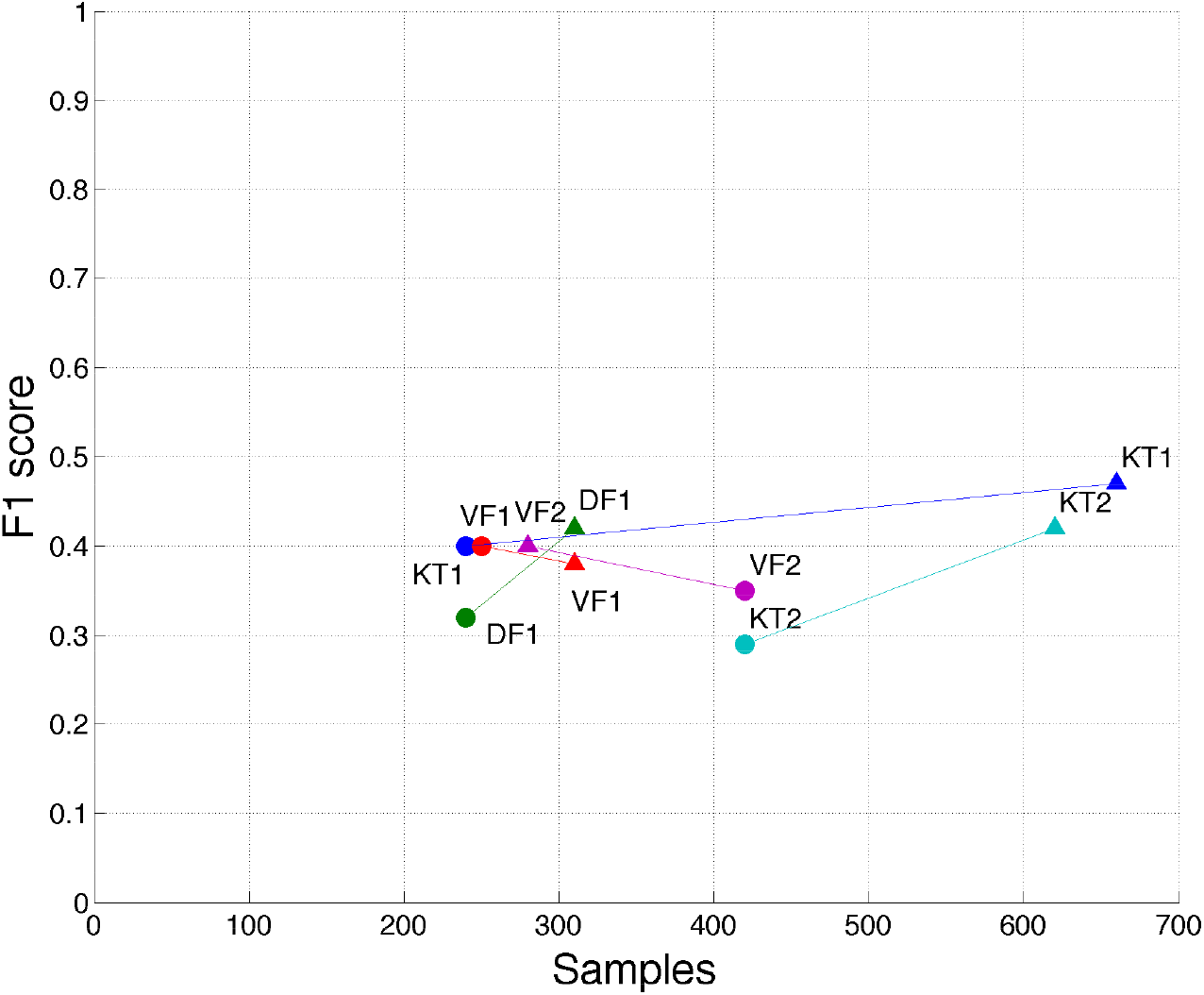
3-class training results using power of thought via traditional (circle) and interactive (triangle) approach. Each test subject is marked with a different color. Horizontal axis shows the number of sample is took to train the model, while the vertical one indicates the performance of the final model on the test run. The experiment continued for as long as test subject felt necessary.

## 5. Discussion

In this work we have proposed, implemented and empirically tested an alternative approach to closed loop feedback BCI training that relies on the visualization of test subject’s mental state space in real time facilitating the interaction with the learning system. In the traditional approach the feedback serves only one purpose – to inform the test subject on his current performance. We expand the information available to the subject by allowing him to see not only that his current actions work poorly (or well), but also provide him with an insight into why a particular set of mental actions might not be a suitable one. We then provide him with an interactive way to experiment with other mental actions until he finds the ones that he can engage consistently and that are distinguishable by the learning system. By the sheer fact of sharing more information with the user we expect our system to achieve better performance. By facilitating the interactive training process we enable the test subject, given enough effort, to reach the desired level of performance.

In addition to the primary benefit described above we find that a few other properties of our approach are beneficial for training BCI systems, namely: the resulting paradigm is personalized to each particular test subject and thus can be tuned better than a one-fits-all paradigm such as motor imagery; system automatically takes care of failed trials and mistakes on the test subject’s side – a subject can rectify a mistake via further interaction with the system, the failed record does not taint the dataset forever, but is gradually phased out by further training; flexibility in training time allows to deviate from strict stimulation schedule and allows the test subject to focus on the most problematic actions, giving them more attention if more attention is needed.

We would like to highlight the choice of the testing paradigm we employed in our work. We find that testing of a general-purpose BCI system must be decoupled from the training in terms of visual cues and protocol. This is necessary to avoid training the subject to simply react to the visual cues and not engage in the corresponding mental activity. By changing the cues we ensure that during the test time the test subject has to invoke the mental activity corresponding to each particular action. Such approach makes the resulting model more robust in the context of real-world applications.

We acknowledge the shortcoming of this study, such as low number of test subjects and a low-end EEG device. This work serves the purpose of initial validation of the concept that allows to plan a larger study, ideally involving intracranial neuroimaging techniques that would have sufficient SNR to make the approach applicable for real-world applications. Rectifying the above-mentioned issues and further exploring the possible topology-preserving dimensionality reduction techniques such as t-SNE [51] and neural networks-based solutions are the primary directions for our future work.

## Acknowledgments

We thank Sven Laur for thorough review of the early version of this work and his suggestions. I.K. and R.V. thank the financial support from the Estonian Research Council through the personal research grant PUT438. This work was supported by the Estonian Centre of Excellence in IT (EXCITE), funded by the European Regional Development Fund.

## Competing interests

The authors declare no competing interests.

